# Device-Derived Social Jet Lag in CPAP-Treated Obstructive Sleep Apnea: Prevalence, Behavioral Signatures, and Age Gradients

**DOI:** 10.64898/2026.06.06.729437

**Authors:** Zachary Yousef, Vedant Ramabadran, Matthew T Scarf, Ioannis P. Androulakis

## Abstract

**Background:** Social jet lag (SJL), the discrepancy timing between work nights and free nights, reflects schedule-related circadian misalignment. Time-stamped CPAP adherence records may provide objective, longitudinal estimates of sleep timing and could augment conventional CPAP reports by adding information on sleep regularity and weekday–weekend misalignment.

**Objectives:** To quantify CPAP-derived SJL in two independent clinical cohorts, characterize its behavioral correlates and age-related patterns, and assess cross-site reproducibility.

**Methods:** We analyzed CPAP-derived sleep timing in patients from Rutgers-RWJ Health (RWJ, N = 1,437) and Hackensack Meridian Health (HMH, N = 1,510) with at least 31 valid nights and at least one valid work night and free night. Mid-sleep on work nights (MSW) and free nights (MSF) was estimated using circular statistics. SJL was defined as the absolute circular difference between MSF and MSW and categorized as none (<1 h), moderate (1–2 h), or severe (≥2 h). Sleep duration, free-night rebound, age-stratified prevalence, and cross-site differences were evaluated using nonparametric and categorical tests.

**Results:** SJL was right-skewed at both sites, with median values below 0.5 h at RWJ and HMH. SJL >1 h was present in 21.2% and 16.4% of patients, respectively; severe SJL occurred in 4.0% and 2.8%. Moderate and severe SJL were associated with shorter work-night sleep and greater free-night rebound, consistent with weekday restriction and weekend compensation. SJL prevalence and variability were highest in younger and middle-aged adults, particularly those aged 26–50 years, and declined markedly after age 65. Core timing phenotypes, including MSW, MSF, and free-night rebound, were highly reproducible across sites despite modest differences in absolute sleep duration and overall SJL prevalence.

**Conclusions:** In CPAP-treated cohorts, SJL is common but usually modest, is associated with weekday sleep restriction and free-night rebound, and declines substantially with age. These findings support the use of routinely collected CPAP data as a scalable, low-burden source of device-anchored circadian screening phenotypes. CPAP-derived SJL may augment standard adherence reports by helping identify patients who warrant further behavioral, circadian, or activity-based assessment.

## Introduction

Social jet lag (SJL) describes the discrepancy between biologically preferred sleep timing and socially imposed schedules (“jet lag without travel”) arising when individuals sleep at systematically different times on work nights versus free nights (Wittmann, Dinich et al. 2006). SJL is commonly quantified as the difference between mid-sleep on work nights (MSW) and mid-sleep on free nights (MSF). Because nights were indexed by sleep-onset date, Sunday–Thursday nights were classified as work nights, corresponding to sleep before Monday–Friday obligations, and Friday–Saturday nights were classified as free nights.Since its initial description, SJL has emerged as a key marker of schedule-related circadian misalignment linked to obesity, metabolic dysfunction, cardiovascular risk, mood disorders, and impaired performance(Wittmann, Dinich et al. 2006, Roenneberg, Allebrandt et al. 2012, Parsons, Moffitt et al. 2015).

SJL is relevant to sleep health because it can represent circadian misalignment even when total sleep time appears preserved. The characteristic behavioral signature is weekday restriction followed by weekend recovery: individuals sleep less and often earlier during work nights, then compensate with longer or later sleep on free nights. While this compensatory pattern may maintain weekly-average sleep duration, evidence suggests that weekend catch-up sleep does not fully reverse the metabolic, inflammatory, and neurobehavioral consequences of weekday sleep restriction (Pejovic, Basta et al. 2013, Depner, Melanson et al. 2019). Thus, SJL captures a form of circadian and behavioral disruption that is distinct from simple sleep deprivation.

Obstructive sleep apnea (OSA) is a common, chronic disease in the United States and worldwide (Senaratna, Perret et al. 2017). The healthcare costs of OSA in the United States are staggering(Wickwire 2021). OSA is due to the intermittent collapse of the upper airway during sleep, which can lead to respiratory pauses, oxygen desaturation, and sleep disruption, among other problems. OSA is highly prevalent, with an estimated 936 million adults aged 30–69 years worldwide affected by mild to severe OSA, including approximately 425 million with moderate to severe disease(Benjafield, Ayas et al. 2019). Continuous positive airway pressure (CPAP) is the first-line treatment for moderate to severe OSA, but real-world adherence remains a major challenge, with approximately 30–50% of patients discontinuing therapy within the first year(Weaver and Grunstein 2008).

The clinical significance of SJL may be especially important in obstructive sleep apnea (OSA), a disorder in which symptoms such as sleepiness often persists even when respiratory events are sufficiently treated (Scharf, Greenberg et al. 2023). In this setting, irregular sleep timing and work–free day differences in sleep behavior may represent an underrecognized contributor to residual symptoms. Thus, examining SJL in CPAP-treated patients provides an opportunity to move beyond respiratory control and duration of use and assess whether routinely collected treatment data can also reveal clinically meaningful circadian and behavioral sleep phenotypes.

In routine clinical practice, CPAP data typically evaluated include the residual apnea–hypopnea index, mask leak, pressure, and duration of use. These measures are essential for evaluating treatment effectiveness, but they provide limited insight into the timing and regularity of CPAP use. A patient may appear adequately treated from a respiratory and adherence standpoint while still experiencing irregular sleep timing, weekday sleep restriction, or weekend compensation that contributes to persistent sleepiness or impaired daytime function.

Time-stamped usage data may be assessed for clinically useful circadian screening phenotypes that could indicate when further self-reported or objective testing is warranted. Because CPAP devices already collect time-stamped usage data longitudinally and passively, they can potentially provide objective estimates of when sleep opportunity occurs, how stable that timing is across the week, and whether work-night and free-night sleep patterns diverge. In this sense, CPAP-derived SJL is not intended to replace actigraphy, sleep diaries, or consumer wearable data. Activity-based approaches can independently capture rest–activity rhythms, movement, naps, non-CPAP sleep, and broader circadian behavior. By contrast, CPAP-derived SJL is a treatment-device–anchored phenotype: it reflects the timing of sleep periods during which the patient used CPAP. Its advantage is clinical accessibility. It can be derived from data already generated during OSA care, without requiring additional devices, patient burden, or separate monitoring protocols.

This distinction provides an important motivation for studying SJL in CPAP cohorts. If reliable timing phenotypes can be extracted from CPAP adherence records, standard CPAP reports could be expanded from “how long was the device used?” to include “when did the patient sleep?” and “how consistent was sleep timing across work and free nights?” Such information could help clinicians differentiate residual sleep-disordered breathing from behavioral or circadian contributors to persistent symptoms, identify patients who may benefit from sleep-schedule counseling, and determine when more detailed assessment with actigraphy or sleep diaries is warranted.

However, CPAP-derived estimates also introduce methodological challenges. Unlike actigraphy or continuous wearable monitoring, CPAP data are adherence-contingent: valid timing data exist only for periods when the device is used. Nights without valid data are unlikely to be missing at random and may cluster by day type, sleep timing, symptom burden, or adherence behavior (Honma, Nohara et al. 2024). CPAP usage timing may also differ from true sleep timing if patients fall asleep before applying the mask, remain awake while using CPAP, remove the mask early, or sleep without the device. Therefore, before CPAP-derived SJL can be used as a clinically interpretable phenotype, its distribution, behavioral signature, and reproducibility across independent clinical cohorts must be characterized.

In this study, we quantify SJL in two large, independent CPAP cohorts, RWJ (N = 1,437) and HMH (N = 1,510), to evaluate whether routinely collected CPAP data can support objective, scalable device-derived circadian screening phenotype in treated OSA. We address three primary questions: (1) What is the distribution, prevalence, and severity of SJL in CPAP-treated patients? (2) Does CPAP-derived SJL show the expected behavioral signature of weekday sleep restriction and free-night rebound? (3) Are CPAP-derived SJL phenotypes reproducible across independent clinical sites? Additionally, we characterize age gradients in SJL to assess whether the well-described decline in SJL with aging is detectable using CPAP-derived timing data.

We hypothesized that: (1) SJL would be prevalent but right-skewed, with most patients showing modest misalignment and a smaller subset showing severe misalignment; (2) moderate and severe SJL would be associated with shorter work-night sleep and compensatory free-night rebound; (3) SJL prevalence would decline with age, particularly after typical retirement age; and (4) core timing phenotypes derived from CPAP usage data would be consistent across sites despite differences in cohort composition or absolute sleep duration.

## Methods

### Study Design and Data Sources

A retrospective cohort study was performed on the population served by the Comprehensive Sleep Disorders Center at RWJ Medical School and the Sleep Center (southern region) at HMH Network. CPAP adherence records on patients seen between 2010-2025 were reviewed. Since this study was done on a clinical population, in keeping with insurance requirements for CPAP therapy, patients with an AHI≥15 or those with an AHI≥5 and a recognized comorbid condition were offered CPAP therapy. De-identified CPAP adherence data was extracted from the ResMed AirView website. The CPAP use times for each 24-hour period were recorded. On the ResMed Airview website, the beginning of a 24-hour period starts at noon. Where there were more than 12 on and off periods of CPAP use per 24 hours, the patient was excluded since accurate data could not be extracted from the Airview website. The institutional review boards at Rutgers Robert Wood Johnson Medical School (Rutgers Biomedical and Health Sciences IRB- Pro2023001290) and HMH Network (HMH Network IRB- Pro2025-0172) approved this study and ethical standards were observed during the study.

### Inclusion Criteria and Analytic Sample

To be included in the SJL analysis, patients were required to have: (1) more than 31 valid nights of CPAP records, and (2) at least one valid work night and valid free night to estimate MSW and MSF. Eight patients in the RWJ cohort and two in the HMH cohort failed to meet completeness criteria and were excluded. The analytic samples comprised 1,437 patients (RWJ) and 1,510 patients (HMH), for a combined total of 2,947 patients.

### Sleep Timing Measures

For each patient, daily CPAP window was approximated by a two-level piecewise constant function, defined by a main usage interval and a background usage level outside that interval (Scharf and Androulakis 2025). The parameters *t*_*on*_ and *t*_*off*_, representing the expected times of CPAP initiation and discontinuation, respectively, were estimated jointly with the corresponding usage levels by minimizing the sum of squared errors between the observed average profile and the fitted piecewise approximation. Thus, *t*_*on*_ and *t*_*off*_ capture the start and end of the patient’s typical daily CPAP-use interval over the analyzed period. In other words they are used as surrogates for sleep onset and wake time(Scharf and Androulakis 2025).

CPAP sleep onset (*t*_*on*_) and sleep offset (*t*_*off*_) were identified using a two-level step function fit via least-squares to the minute-by-minute binary mask of CPAP usage (Scharf, Androulakis, 2025). For each patient, CPAP timing was estimated from the patient’s average daily usage profile over the analyzed recording period, rather than by fitting separate onset and offset times for each individual night; the resulting *t*_*on*_ and *t*_*off*_ therefore represent the patient’s typical CPAP-use interval for the relevant day type.Nightly mid-sleep (MS) was computed as the circular midpoint between *t*_*on*_ and *t*_*off*_.

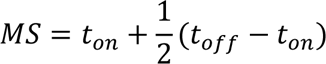

Because clock times wrap at midnight, circular statistics were used throughout. For each patient, MSW was the circular mean of nightly mid-sleep on work nights (Sunday–Thursday), and MSF was the circular mean of nightly mid-sleep on free/weekend nights (Friday–Saturday). SJL was defined as the circular difference: *SJL* = *MSF* − *MSW*. A positive SJL reflecting delayed free-night sleep(Wittmann, Dinich et al. 2006, Jankowski 2017, Roenneberg, Pilz et al. 2019). Severity categories were defined as: no SJL (SJL < 1h), moderate (1 ℎ ≤ *SJL* < 2 ℎ), and severe (*SJL* ≥ 2 ℎ), following the thresholds established in the literature(Wittmann, Dinich et al. 2006).

The signed circular difference *MSF* − *MSW*was calculated to describe the direction of the work-night to free-night timing shift, with positive values indicating later free-night mid-sleep and negative values indicating earlier free-night mid-sleep. Negative signed differences were rare in both cohorts and, upon inspection, were associated with fragmented or irregular CPAP-use patterns suggestive of unreliable timing estimation. Because these records were more consistent with poor-quality or nonrepresentative device-use data than with a stable earlier free-night sleep phenotype, patients with negative signed SJL were excluded from the primary analytic dataset. SJL severity was then categorized using the remaining positive signed differences, with thresholds of <1 h, 1–2 h, and ≥2 h.

### Sleep Duration and Weekend Rebound

Sleep duration per night was computed as the circular distance between *t*_*on*_ and *t*_*off*_. For each patient, mean sleep duration was calculated separately for work nights and free nights. Free-night rebound was defined as the per-patient difference: mean(free-night sleep duration) − mean(work-night sleep duration). A positive rebound indicates compensatory sleep extension on Friday and Saturday relative to Sunday–Thursday. Rebound significance within SJL categories was tested using Wilcoxon signed-rank tests (paired, non-parametric).

### Age-Stratified Analyses

Patients were stratified into eight age groups: 19–25, 26–35, 36–50, 51–65, 66–70, 71–75, 76–80, and ≥80 years. The broader bins at younger ages (19–25, 26–35, 36–50) reflect lower OSA prevalence and smaller sample sizes in younger populations(Senaratna, Perret et al. 2017) the transition to narrower 5-year bins at age 66 follows precedent from (Prigent, Blanloeil et al. 2023) who identified a clinically significant adherence inflection after age 80 that broader bins would obscure. Prevalence of SJL > 1 h was estimated within each stratum using Wilson 95% confidence intervals, which provide reliable coverage even with small samples or extreme proportions because they do not rely on normal approximation(Brown, Cai et al. 2001). Association between age and SJL prevalence was tested using chi-square tests of independence (Bewick, Cheek et al. 2004) and Kruskal–Wallis H tests on continuous SJL. Pairwise comparisons used Fisher’s exact tests with Benjamini–Hochberg false discovery rate (FDR) correction(Benjamini and Hochberg 1995).

### Cross-Site Comparison

Ten measures were compared between RWJ and HMH: SJL distribution, work-night sleep duration, free-night sleep duration, free-night rebound, weekly-average sleep, valid nights, MSW, MSF, SJL > 1 h prevalence, and severity category distribution. Continuous comparisons used Mann-Whitney U tests; categorical comparisons used chi-square tests. Statistically significant differences were defined as p ≤ 0.05 (α = 0.05).

### Statistical Analysis

All analyses were performed at the patient level. Continuous variables are summarized as mean ± SD or median [IQR], selected using the criterion: for positive-valued variables where the mean/SD ratio falls below 2, median [IQR] is preferred as the more robust summary of right-skewed distributions(Mansournia and Nazemipour 2024). Categorical variables are reported as counts (%). Statistical significance was set at α = 0.05, two-sided. All analyses were conducted in Python using circular statistics (CircStat library) for timing variables and SciPy for hypothesis tests.

## Results

### Cohort Characteristics

**Table 1** presents cohort characteristics for both sites. The RWJ analytic cohort comprised 1,437 patients (mean age 59.5 ± 13.2 years; median age 60.8 years [IQR 52.1–68.5]). Sex was recorded for only 14.7% of RWJ patients (9.7% male, 5.0% female), reflecting incomplete demographic capture at this site. Patients contributed a median of 468 valid nights [IQR 161–874], with median work nights of 338 [117–631] and free nights of 126 [42–244]. The age distribution was concentrated in the 51–65 group (42.9%), **Table 2**.

**Table 1:** Cohort Characteristics — RWJ vs. HMH. Data presented as mean ± SD, median [IQR], or count (%). IQR: interquartile range. Valid nights include all nights meeting quality criteria; work nights: Sunday–Thursday; free nights: Friday–Saturday.

| Variable | RWJ (N = 1,414) | HMH (N = 1,496) |
| --- | --- | --- |
| Age, mean ± SD | 59.5 ± 13.2 yrs | 61.9 ± 13.5 yrs |
| Age, median [IQR] | 60.8 [52.1–68.5] | 62.7 [54.6–71.5] |
| Sex: Male | 136 (9.6%) | 317 (21.2%) |
| Sex: Female | 71 (5.0%) | 188 (12.6%) |
| Sex: Not recorded | 1,207 (85.4%) | 991 (66.2%) |
| Valid nights, median [IQR] | 474 [166-890] | 478 [171-874] |
| Valid work nights, median [IQR] | 343 [120-638] | 341 [123–631] |
| Valid free nights, median [IQR] | 128 [43–244] | 131 [46–246] |

**Table 2:** Age Group Distribution — RWJ vs. HMH. Age stratification follows(Prigent, Blanloeil et al. 2023) for narrower 5-year bins from age 66 onward. Combined column reflects the full analytic cohort (N = 2,910).

| Age Group | RWJ N (%) | HMH N (%) |
| --- | --- | --- |
| 19–25 | 11 (0.8%) | 6 (0.4%) |
| 26–35 | 71 (5.0%) | 54 (3.6%) |
| 36–50 | 247 (17.5%) | 218 (14.6%) |
| 51–65 | 604 (42.7%) | 608 (40.6%) |
| 66–70 | 232 (16.4%) | 217 (14.5%) |
| 71–75 | 118 (8.3%) | 173 (11.6%) |
| 76–80 | 79 (5.6%) | 115 (7.7%) |
| 80+ | 52 (3.7%) | 105 (7.0%) |

The HMH analytic cohort comprised 1,496 patients (mean age 61.9 ± 13.5 years; median age 62.7 [54.6–71.5]). Sex recording was substantially better at this site (34.0% recorded: 21.5% male, 12.5% female). The HMH cohort was slightly older, with a larger proportion of patients aged 71 and older (26.2% vs. 17.3% at RWJ). Valid night counts were similar across sites. **Table 3** presents the combined age distribution across sites.

**Table 3:** Cross-Site Statistical Comparison — RWJ vs. HMH. All continuous values are median [IQR]. Mann-Whitney U: non-parametric test comparing two independent distributions. Chi-square: comparing categorical frequency distributions. * Statistically significant (p < 0.05). NS: not significant. SIMILAR: p ≥ 0.05; DISSIMILAR: p < 0.05. MSW: mid-sleep on work nights; MSF: mid-sleep on free nights.

| Measure | RWJ | HMH | p-value |
| --- | --- | --- | --- |
| SJL (h), median [IQR] | 0.46 [0.20–0.86] | 0.41 [0.18–0.79] | $p = 0.01^*$ |
| Work-night sleep (h) | 6.26 [5.22–7.20] | 6.61 [5.40–7.52] | $p < .001^*$ |
| Free-night sleep (h) | 6.43 [5.27–7.48] | 6.75 [5.51–7.76] | $p < .001^*$ |
| Free-night rebound (h) | 0.09 [−0.14–0.38] | 0.10 [−0.11–0.36] | $p = 0.46$ |
| Weekly avg sleep (h) | 6.32 [5.23–7.27] | 6.65 [5.43–7.58] | $p < .001^*$ |
| Valid nights | 474 [166–890] | 477 [171–874] | $p = 0.89$ |
| MSW (h) | 14.16 [13.27–15.20] | 14.21 [13.29–15.13] | $p = 0.68$ |
| MSF (h) | 14.75 [13.76–15.74] | 14.72 [13.86–15.61] | $p = 0.68$ |
| SJL > 1 h prevalence | 19.9% (282/1414) | 15.7% (235/1496) | $p = 0.003^*$ |
| Severity distribution | 1132 / 234 / 48 | 1261 / 198 / 37 | $p = 0.01^*$ |

### SJL Distribution and Severity

At both sites, SJL exhibited a strongly right-skewed distribution: the majority of patients had small work–free timing differences, with a long tail toward higher SJL. **Figure 1** shows the distribution of social jetlag (SJL) at RWJ and HMH, with patients classified into no SJL (<1 h), moderate SJL (1–2 h), and severe SJL (≥2 h). At both sites, the distribution was strongly right-skewed, with most patients falling below the 1-hour threshold. At RWJ, 1,132 of 1,415 patients were classified as having no SJL, while 234 had moderate SJL and 49 had severe SJL. The overall median SJL was 0.46 h, with a mean of 0.64 h and SD of 0.72 h. At HMH, a similar distribution was observed: 1,262 of 1,497 patients had no SJL, 198 had moderate SJL, and 37 had severe SJL. The median SJL was slightly lower than at RWJ, at 0.41 h, with a mean of 0.57 h and SD of 0.58 h.

**Figure 1:**
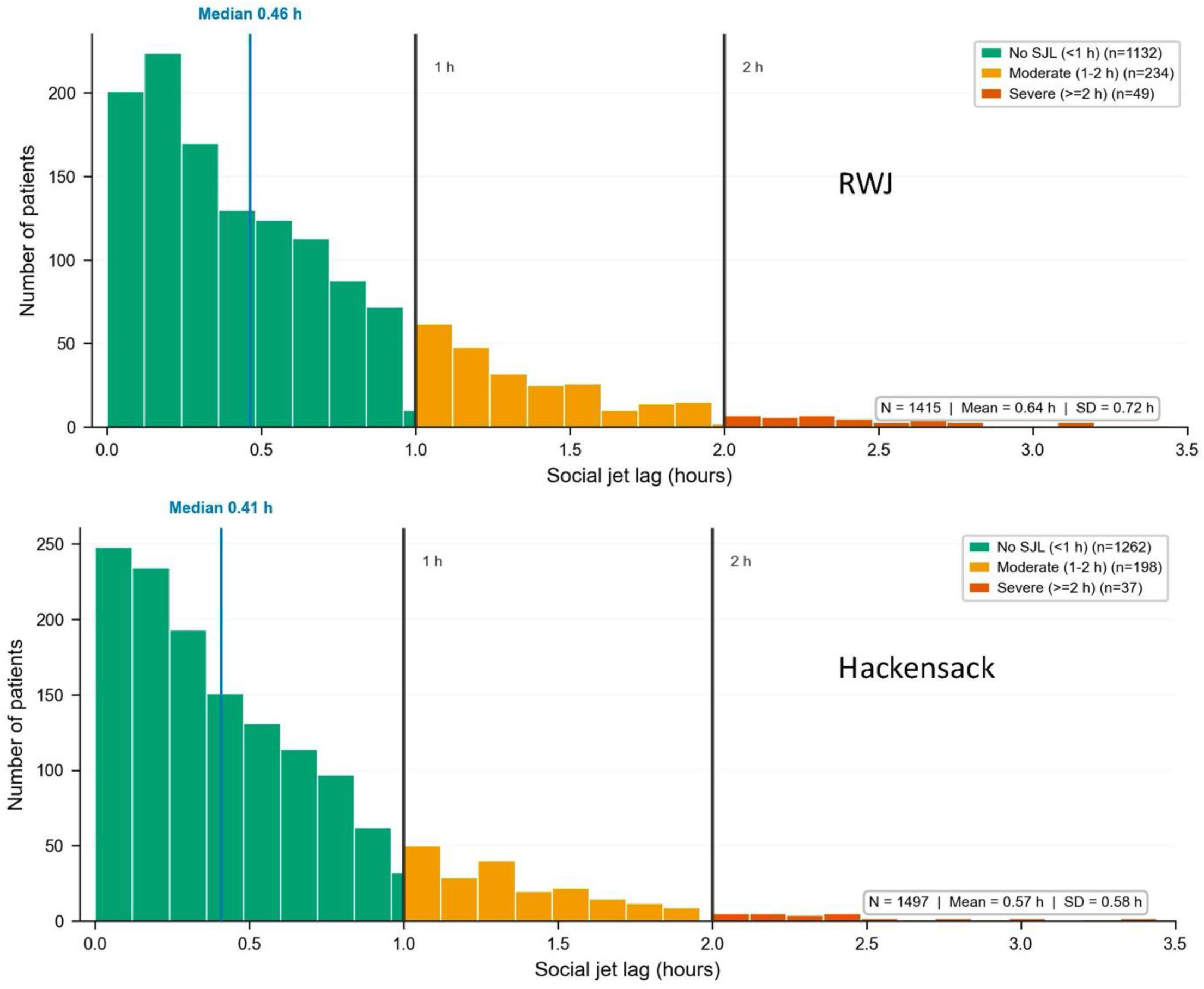
Distribution of social jetlag at RWJ and HMH. Histograms show the distribution of social jetlag (SJL, hours) in patients from RWJ and HMH. Vertical reference lines indicate the predefined thresholds for moderate SJL (≥1 h) and severe SJL (≥2 h), and the blue line indicates the cohort median. Bars are color-coded by SJL category: no SJL (<1 h), moderate SJL (1–2 h), and severe SJL (≥2 h). Both cohorts showed right-skewed distributions, with most patients having SJL <1 h and a smaller subset demonstrating moderate or severe SJL.

Signed SJL values were predominantly positive, indicating later free-night mid-sleep among patients with measurable SJL. Negative signed values were rare and were generally associated with more fragmented or irregular CPAP-use patterns, suggesting that they were more likely to reflect lower-quality timing estimates than a consistent biological phenotype.

Overall, these distributions indicate that although most patients had relatively low SJL, a substantial minority exceeded clinically interpretable thresholds. Moderate SJL was present in approximately 16.5% of RWJ patients and 13.2% of HMH patients, while severe SJL was observed in approximately 3.5% and 2.5%, respectively. The similar right-skewed pattern across both sites supports the reproducibility of the SJL distribution and suggests that meaningful sleep-timing misalignment is present in a nontrivial subset of patients.

### Mid-sleep patterns and SJL

Across both sites, weekly mid-sleep profiles showed a similar overall structure, with clear separation by SJL severity, but also highlighted important heterogeneity within the severe group, **Figure 2** illustrates day-of-week variation in mid-sleep timing by SJL severity group at RWJ and HMH.

**Figure 2:**
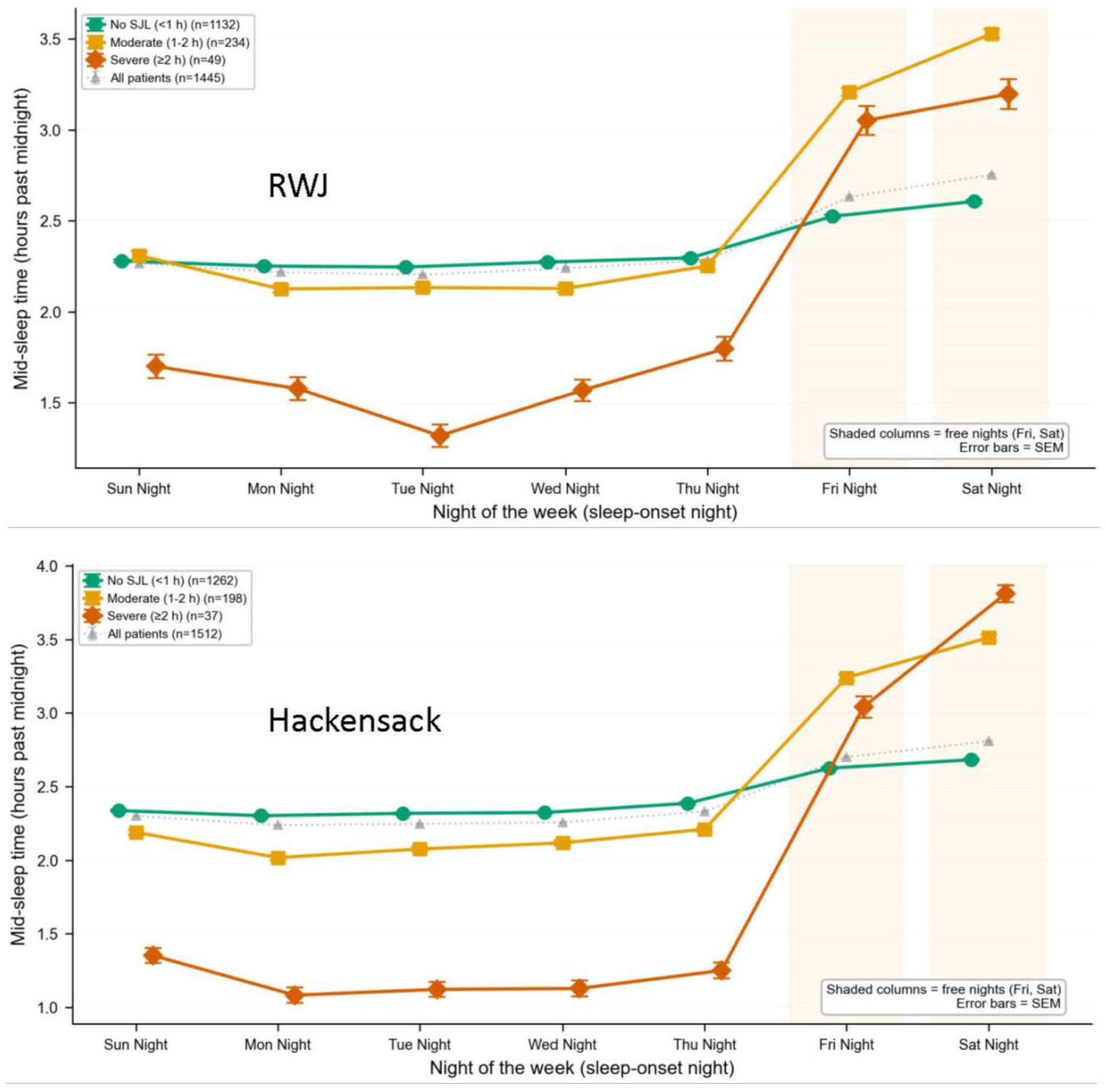
Day-of-week variation in mid-sleep timing by SJL category. Mean mid-sleep time, expressed as hours past midnight, is shown by sleep-onset night and SJL category for RWJ and HMH. Shaded regions indicate free nights (Friday and Saturday), and error bars represent SEM. Across both sites, patients with no SJL showed relatively stable mid-sleep timing across the week, whereas patients with moderate and severe SJL showed marked delays on free nights. The severe-SJL group exhibited the largest weekday–weekend shift, reflecting earlier/constrained weekday sleep timing and substantially delayed free-night sleep timing.

At both sites, participants with no SJL demonstrated relatively stable mid-sleep timing across the week, with only a small delay on free nights. At RWJ, the no-SJL group remained close to 2.25–2.30 hours past midnight from Sunday through Thursday, increasing modestly to approximately 2.5–2.6 hours on Friday and Saturday nights. At HMH, a similar pattern was observed, with weekday mid-sleep times near 2.3–2.4 hours and free-night values increasing slightly to approximately 2.6–2.7 hours. By contrast, participants with moderate or severe SJL showed a much larger weekday-to-weekend displacement in sleep timing. In the moderate-SJL group, mid-sleep times were generally around 2.0–2.3 hours past midnight during weekday nights, but shifted later on free nights, reaching approximately 3.2–3.5 hours past midnight by Friday and Saturday. The severe-SJL group showed the largest contrast: weekday mid-sleep occurred substantially earlier than in the other groups, especially at HMH, where values were approximately 1.1–1.3 hours past midnight from Monday through Thursday, followed by a sharp weekend delay to approximately 3.0 hours on Friday and 3.8 hours on Saturday. A comparable pattern was observed at RWJ, where severe-SJL mid-sleep shifted from approximately 1.3–1.8 hours during the weekday interval to approximately 3.0–3.2 hours on free nights.

Overall, the figure demonstrates that higher SJL severity reflects a widening separation between weekday and free-night sleep timing. The pattern was consistent across both sites and suggests that severe SJL is driven by a combination of relatively early/constrained weekday sleep timing and substantially delayed sleep timing on free nights. The results indicate that increasing SJL severity is associated not only with later sleep timing on free nights, but also with relatively earlier mid-sleep timing during weekday nights, especially in the severe-SJL group. This pattern was highly consistent across the two sites and suggests that greater SJL reflects a larger discrepancy between constrained weekday sleep schedules and delayed free-night sleep timing.

### Sleep Duration Signatures: Weekday Restriction and Weekend Rebound

Across both cohorts, SJL category was associated with a characteristic pattern of weekday restriction and free-night rebound. **Figure 3** summarizes sleep duration patterns by SJL category and site, comparing work nights (Sunday–Thursday) with free nights (Friday–Saturday). At RWJ, mean sleep duration was modestly longer on free nights than work nights across all SJL groups. Patients with no SJL slept 6.26 h on work nights and 6.33 h on free nights, corresponding to a small but significant free-night rebound of +0.07 h (p < .001). The moderate-SJL group showed a larger increase from 5.90 h on work nights to 6.29 h on free nights, with a rebound of +0.40 h (p < .001). Patients with severe SJL had the shortest work-night sleep duration, averaging 5.70 h, and increased to 6.04 h on free nights, corresponding to a rebound of +0.34 h (p = .016).

**Figure 3:**
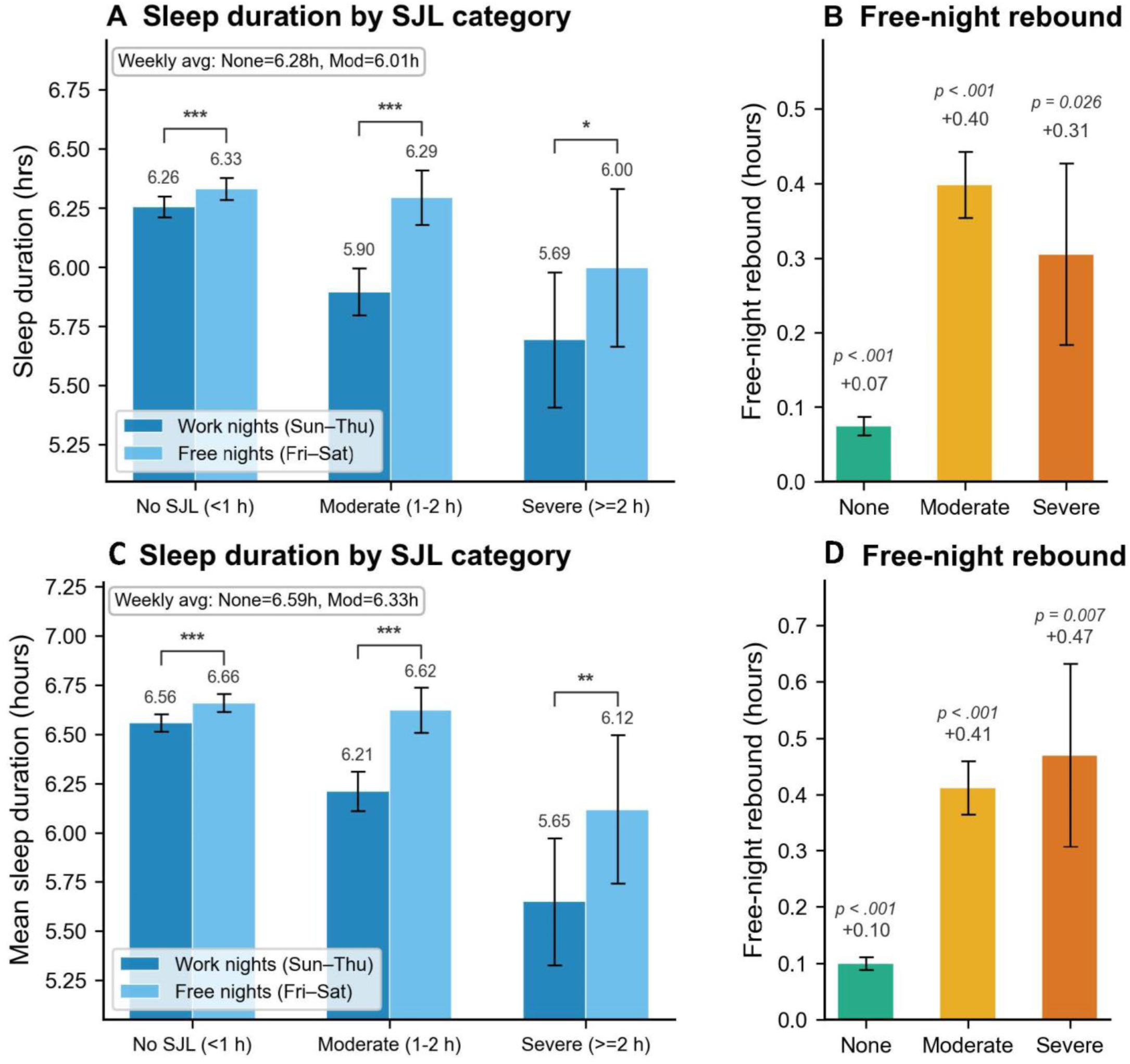
Sleep duration and free-night rebound by SJL category. Panels A and C show mean sleep duration on work nights (Sunday–Thursday) and free nights (Friday–Saturday) by SJL category for RWJ and HMH, respectively. Panels B and D show the corresponding free-night rebound, calculated as the difference between free-night and work-night sleep duration. Error bars represent SEM. Across both sites, sleep duration increased significantly on free nights in all SJL groups, with larger rebounds among patients with moderate or severe SJL than among those with no SJL. Asterisks indicate statistical significance (*p < .05, **p < .01, ***p < .001).

A similar pattern was observed at HMH. Patients with no SJL slept 6.56 h on work nights and 6.66 h on free nights, yielding a small but significant rebound of +0.10 h (p < .001). In the moderate-SJL group, sleep duration increased from 6.21 h on work nights to 6.62 h on free nights, corresponding to a +0.41 h rebound (p < .001). The severe-SJL group again showed the shortest work-night sleep duration, averaging 5.65 h, with an increase to 6.12 h on free nights and a rebound of +0.47 h (p = .007).

Overall, these findings indicate that increasing SJL severity is associated with shorter sleep duration on work nights and a greater compensatory increase in sleep duration on free nights. This pattern was consistent across both sites, supporting the interpretation that SJL reflects not only a shift in sleep timing but also a weekday–weekend imbalance in sleep opportunity or sleep need.

### Age-Stratified SJL Distributions and Mid-Sleep Timing

**Figure 4** shows the prevalence of SJL >1 h across age groups at RWJ and HMH. At both sites, SJL prevalence varied strongly by age, with the highest rates observed in younger and middle-aged adults and substantially lower rates among older adults. At RWJ, prevalence increased from approximately 18% in the 19–25 age group to 32–33% in the 26–35 and 36–50 age groups, then declined progressively to approximately 22% in ages 51–65, 13% in ages 66–70, 7–8% in ages 71–75, and below 5% in patients older than 75 years. HMH showed a similar age-dependent pattern, with SJL prevalence of approximately 17% in ages 19–25, 24% in ages 26–35, 31% in ages 36–50, and 19% in ages 51–65, followed by lower prevalence in older age groups, generally below 10% after age 65.

**Figure 4:**
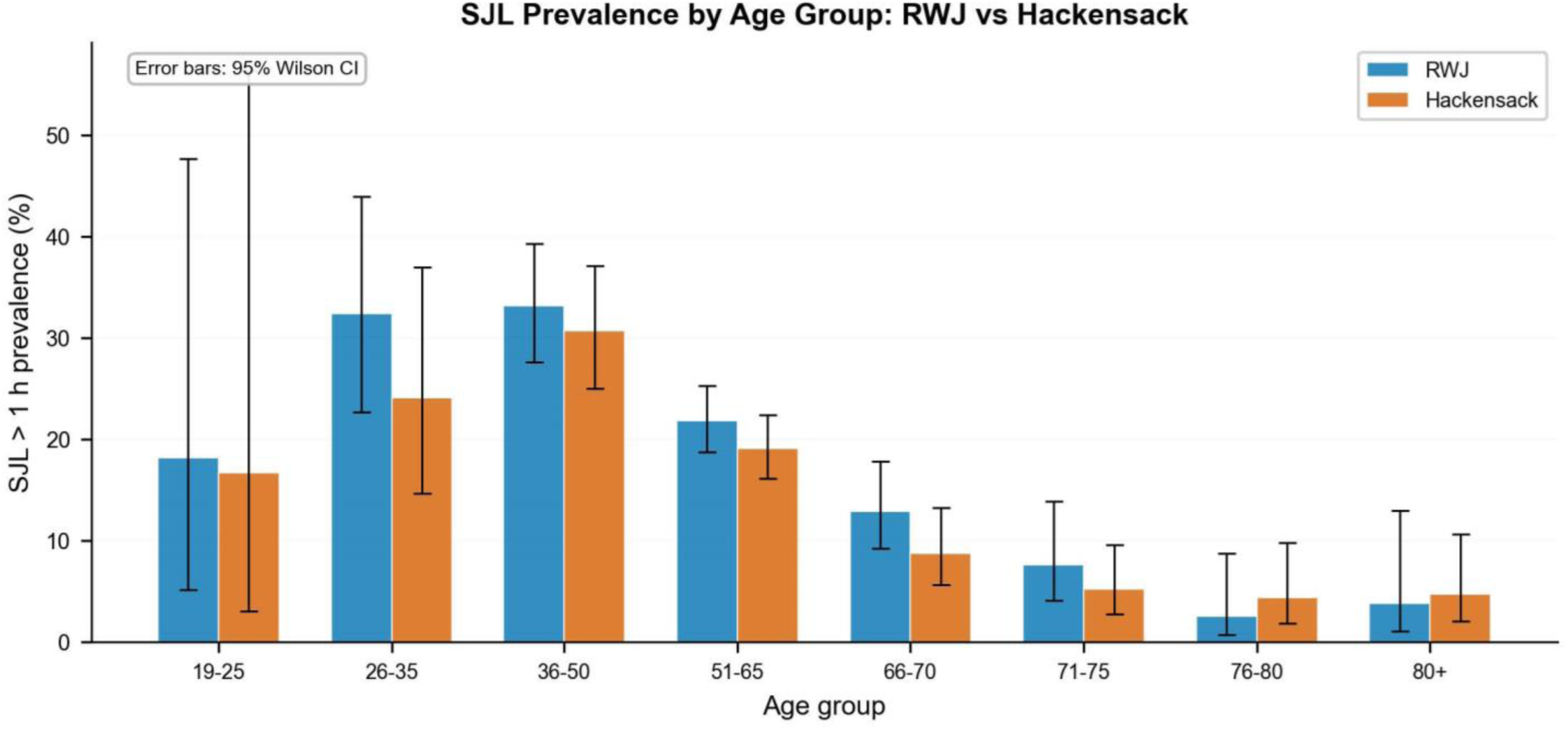
Prevalence of clinically meaningful social jetlag by age group. Bar plots show the prevalence of SJL >1 h across age groups at RWJ and HMH. Error bars represent 95% Wilson confidence intervals. In both cohorts, SJL prevalence was highest among younger and middle-aged adults, particularly those between 26 and 50 years of age, and declined substantially with advancing age. The broad consistency between sites supports the reproducibility of the age-dependent pattern.

Overall, these findings indicate that clinically meaningful SJL is most common among adults younger than 65 years, particularly those between 26 and 50 years of age, and becomes progressively less prevalent with advancing age. The pattern was broadly consistent across the two sites, supporting the reproducibility of the age association. The wider confidence intervals in the youngest and oldest age groups reflect smaller sample sizes in those strata and suggest that estimates for these groups should be interpreted with greater caution.

**Figure 5** further presents mid-sleep plots show that this age pattern reflects differences in both work-night mid-sleep (MSW) and free-night mid-sleep (MSF). Younger adults had the latest mid-sleep times on both work and free nights, with the 19–25 group showing mid-sleep times around 3.5–4.5 hours past midnight. Across subsequent age groups, both MSW and MSF shifted earlier, particularly through midlife, and the separation between free-night and work-night timing generally narrowed with older age. This narrowing is consistent with the lower SJL observed in older patients. Overall, the figure indicates that SJL is most pronounced in younger and middle-aged adults because free-night sleep timing remains delayed relative to work-night timing, whereas older adults show earlier and more closely aligned sleep timing across the week.

**Figure 5:**
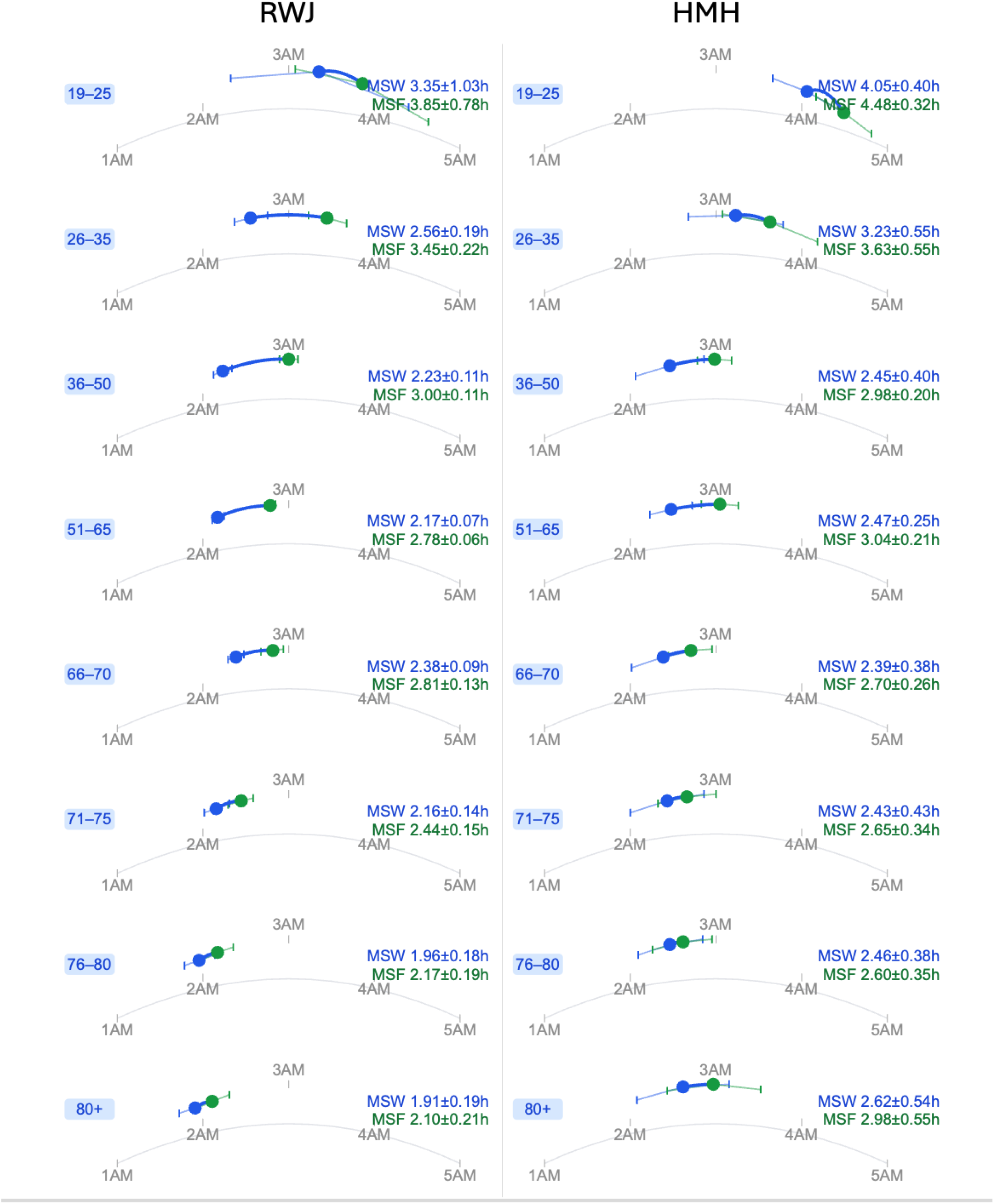
Age-related patterns in social jetlag and mid-sleep timing. Stacked arc plots of work-night mid-sleep (MSW, blue) and free-night mid-sleep (MSF, green) by age group, showing later timing and greater MSW–MSF separation in younger adults, consistent with greater social jet lag.

### Cross-Site Comparison: RWJ vs. HMH

**Table 3** presents the full cross-site statistical comparison across 10 measures. Several core timing measures showed minimal between-site differences, including MSW, MSF, and free-night rebound.: free-night rebound (RWJ: 0.09 h vs. HMH: 0.11 h; p = 0.297), valid night count (468 vs. 476; p = 0.913), MSW (14.17 vs. 14.22 h; p = 0.771), and MSF (14.74 h at both sites; p = 0.921). Six measures were statistically dissimilar: SJL magnitude (median difference 0.05 h; p = 0.007), work-night sleep duration (6.25 vs. 6.60 h; p < .001), free-night sleep duration (6.42 vs. 6.75 h; p < .001), weekly-average sleep (6.30 vs. 6.65 h; p < .001), SJL > 1 h prevalence (21.2% vs. 16.4%; p = 0.001), and severity distribution (p = 0.004).

Critically, the statistically similar measures include the core SJL timing phenotypes: MSW, MSF, and free-night rebound. The difference in median SJL between sites, while statistically significant due to the large sample sizes, is only 0.05 h (∼3 minutes)—likely of limited clinical significance. The larger differences in absolute sleep duration (∼0.30–0.35 h) likely reflect the older age profile of the HMH cohort, which has proportionally more patients in the 71+ age groups where work-night sleep duration is longer. The consistent pattern of age-related SJL decline across both independent sites (chi-square p < .001 at both sites) substantially strengthens external validity.

## Discussion

In this study of two independent CPAP-treated cohorts totaling 2,947 patients, we used objective, longitudinal device-derived timing data to quantify social jet lag and characterize its behavioral and demographic correlates. Four principal findings emerged. First, SJL was common but highly right-skewed: most patients had modest work–free differences in mid-sleep timing, whereas a clinically meaningful subset exceeded the 1-hour threshold and a smaller subgroup exceeded 2 hours. Second, SJL showed a consistent behavioral signature across both sites, with moderate and severe SJL associated with shorter work-night sleep and greater free-night rebound, consistent with weekday restriction and weekend compensation. Third, SJL was strongly age-dependent, with the highest prevalence and variability in younger and middle-aged adults and a marked decline after age 65. Finally, the core timing phenotypes: MSW, MSF, and free-night rebound, were reproducible across RWJ and HMH despite modest site differences in absolute sleep duration and overall SJL prevalence. These findings suggest that CPAP-derived SJL captures structured differences in sleep timing rather than random night-to-night variation. The observed pattern links greater SJL to both timing misalignment and sleep-duration imbalance, while the age gradient is consistent with reduced social scheduling constraints and greater regularity of sleep timing in older adults.

Together, these findings support the feasibility of using CPAP adherence data as a scalable platform for device-derived circadian screening phenotype in clinical sleep populations. Unlike questionnaire-based measures, CPAP records provide repeated, objective, night-by-night estimates over long observation windows. This makes it possible to quantify not only average sleep timing, but also the degree to which timing differs between work nights and free nights. The cross-site reproducibility observed here is particularly important because it suggests that the main SJL phenotypes are not an artifact of a single center, data extraction workflow, or local patient population.

### Interpretation of the SJL Distribution

The distribution of SJL in both cohorts was dominated by values below 1 hour, but with a long right tail. This pattern is consistent with SJL as a heterogeneous behavioral phenotype rather than a uniformly present feature of CPAP-treated OSA. In practical terms, most patients using CPAP did not demonstrate large differences between work-night and free-night mid-sleep timing. However, approximately one in five RWJ patients and one in six HMH patients had SJL greater than 1 hour, indicating that meaningful schedule-related misalignment is present in a substantial minority of patients. The severe group was comparatively small, but its timing profile was distinctive and consistent across sites.

The modest but statistically significant site difference in median SJL should be interpreted cautiously. The absolute difference between sites was approximately 0.05 hours, or about 3 minutes, which is unlikely to be clinically meaningful despite statistical significance in a large sample. In contrast, the similar MSW, MSF, and rebound estimates across sites provide stronger evidence that the underlying timing structure was shared. The observed differences in sleep duration and SJL prevalence may reflect demographic composition, particularly the older age distribution at HMH, rather than true differences in circadian behavior between clinical centers.

### Weekday Restriction and Free-Night Rebound

The sleep-duration analyses help clarify what SJL represents behaviorally in these CPAP cohorts. Patients with moderate and severe SJL slept less on work nights than patients with no SJL and showed larger increases in sleep duration on free nights. This pattern is consistent with weekday restriction followed by partial weekend compensation. Importantly, this rebound occurred alongside delayed free-night mid-sleep timing, indicating that SJL reflects both a timing shift and a sleep-duration imbalance. Thus, SJL should not be interpreted simply as a later weekend schedule; it also appears to mark a pattern of accumulated weekday sleep deficit or constrained sleep opportunity.

The severe-SJL group showed particularly large weekday-weekend timing contrasts. In both cohorts, severe SJL was characterized not only by delayed free-night mid-sleep, but also by relatively early work-night mid-sleep, especially at HMH. This pattern suggests that severe SJL may arise when social or occupational constraints compress and advance sleep timing during work nights, followed by later and longer sleep on free nights. However, this interpretation should be balanced against the measurement limitations of CPAP data: early mask removal or variable adherence could also shift apparent work-night mid-sleep earlier. The consistency of the pattern across sites argues that the signal is meaningful, but future studies incorporating actigraphy or sleep diaries would help distinguish true sleep timing from CPAP-use timing.

### Age Gradient in SJL

Age was one of the strongest correlates of SJL. Clinically meaningful SJL was most prevalent among adults aged 26-50 years and declined progressively in older age groups, particularly after age 65. The age-stratified distribution plots showed that this decline reflected both lower median SJL and reduced between-patient variability. Younger and middle-aged adults had wider SJL distributions and larger separation between work-night and free-night mid-sleep timing, whereas older adults showed more stable and more closely aligned sleep timing across the week.

Several mechanisms may contribute to this pattern. Younger and middle-aged adults are more likely to experience work, family, and social constraints that impose different schedules on weekdays and weekends. In older adults, retirement or reduced work-schedule rigidity may lessen the distinction between work nights and free nights. Age-related changes in circadian phase, sleep timing, and sleep regularity may also contribute. The present data cannot separate retirement status from biological aging or cohort effects, but the replication of the age gradient at two independent sites strengthens the conclusion that CPAP-derived SJL captures a meaningful life-course pattern.

### Clinical Implications

CPAP-derived SJL could augment standard CPAP reports by adding timing regularity and weekday–weekend misalignment information to conventional adherence and respiratory metrics. Current clinical reports primarily describe whether and how effectively the device was used; SJL phenotyping adds information about when CPAP-supported sleep occurs and whether that timing differs across the week.

SJL phenotyping in CPAP cohorts has several potential clinical applications. First, it may help identify patients with persistent sleepiness, or impaired daytime function despite apparently adequate CPAP treatment. In such patients, residual symptoms may reflect schedule-related circadian misalignment, insufficient work-night sleep, or weekend compensation rather than residual sleep-disordered breathing alone. Incorporating SJL metrics into CPAP review could therefore broaden the clinical interpretation of adherence reports from hours of use alone to include when sleep is occurring and how stable that timing is across the week.

Second, the association of moderate and severe SJL with shorter work-night sleep duration points to a modifiable behavioral target. Counseling could focus on extending work-night sleep opportunity, reducing large weekend delays in sleep timing, maintaining a more stable wake time, and identifying occupational or family constraints that drive weekday restriction. These interventions may be especially relevant for younger and middle-aged CPAP users, in whom SJL prevalence was highest. Because CPAP therapy addresses respiratory events but does not necessarily correct irregular sleep schedules, combining PAP optimization with circadian and behavioral sleep counseling may improve patient-centered outcomes.

Third, device-derived SJL may provide a scalable screening marker for clinical workflows and future interventional studies. Patients with SJL greater than 1 hour could be flagged for targeted assessment of sleep timing, chronotype, shift work, residual sleepiness, and sleep sufficiency. At the population level, CPAP-derived timing phenotypes may also help identify subgroups at risk for poor adherence or persistent symptoms, supporting more personalized sleep medicine.

### Limitations

Several limitations should be considered. CPAP-derived SJL is a device-anchored estimate of sleep timing, not a direct measure of endogenous circadian phase or a complete rest–activity rhythm. Unlike actigraphy, sleep diaries, or other wearable activity measures, CPAP data capture only periods when the patient is using the device. Thus, CPAP-derived timing phenotypes are clinically accessible and scalable, but they are inherently conditional on treatment use. Directionality estimates may be particularly sensitive to CPAP-use quality; in this dataset, rare negative signed SJL values were often associated with fragmented or irregular usage patterns, reinforcing the decision to focus primary analyses on absolute SJL magnitude.

CPAP-derived sleep estimates are adherence-contingent. Valid timing data exist only for nights when the device is used, and missingness is unlikely to be random. Patients with irregular sleep-wake patterns may have lower adherence, more fragmented device use, or selective non-use on certain nights, meaning that those with the greatest circadian irregularity could be underrepresented. The requirement for at least 31 valid nights and at least one work night and one free night was necessary for stable estimation, but it may also preferentially exclude patients with the most inconsistent CPAP use. As a result, the observed SJL estimates may be conservative.

CPAP use is an imperfect proxy for sleep. Patients may fall asleep before applying the mask, remain awake after starting CPAP, remove the mask before final awakening, use CPAP during naps, or sleep for part of the night without CPAP. These behaviors could bias estimates of sleep onset, sleep offset, sleep duration, and mid-sleep timing. Early mask removal is particularly relevant because it could artificially shorten estimated sleep duration and advance apparent mid-sleep, especially on work nights. Validation against actigraphy, sleep diaries, or polysomnography-derived timing would strengthen interpretation of CPAP-derived SJL and help determine when CPAP-derived estimates are sufficiently accurate for clinical use.

The classification of Sunday–Thursday as work nights and Friday–Saturday as free nights is a simplifying assumption. This convention may not apply to shift workers, healthcare workers, retirees, unemployed individuals, caregivers, or patients with non-standard schedules. This limitation is especially important in older age groups, where calendar-based work/free labels may not correspond to actual social obligations. Future studies should infer individualized work–free structure where possible, incorporate employment status and work schedule information, or combine CPAP data with sleep diaries or actigraphy to better define socially constrained and unconstrained nights.

Available clinical metadata were limited. Sex was incompletely recorded at both sites, especially at RWJ, precluding reliable sex-stratified analyses. We also lacked information on race and ethnicity, employment status, shift work, comorbidities, medications, residual AHI, sleepiness scores, insomnia symptoms, chronotype, socioeconomic factors, and daytime activity patterns. These variables may influence both CPAP adherence and sleep timing and may partially explain between-site differences in sleep duration and SJL prevalence. The observational design also precludes causal inference; the data show associations among age, SJL, sleep duration, and rebound, but cannot determine whether SJL causes persistent symptoms, poor adherence, or adverse clinical outcomes.

Finally, the severe-SJL subgroup was relatively small, particularly at HMH, which increases uncertainty around estimates for this category. Circular averaging provides a robust way to summarize clock times, but it may obscure multimodal, highly irregular, or fragmented sleep schedules. Therefore, these results should be interpreted as population-level evidence of reproducible timing patterns, not as a complete characterization of all forms of circadian disruption in CPAP-treated OSA. Collectively, these limitations suggest that CPAP-derived SJL is best viewed as a conservative, adherence-aware screening phenotype that can augment conventional CPAP reports but requires further validation against independent activity-based and clinical outcome measures.

## Conclusions

In two large, independent CPAP-treated cohorts totaling nearly 3,000 patients, social jet lag was common but typically modest, with right-skewed distributions and median values below 0.5 hours at both sites. Approximately one in five patients had SJL greater than 1 hour, and a smaller subset had severe SJL of at least 2 hours. The reproducibility of these findings across RWJ and HMH supports the feasibility of using routinely collected CPAP data for objective, large-scale device-derived circadian screening phenotype in treated obstructive sleep apnea populations. The behavioral signature of SJL was consistent across sites. Moderate or severe SJL was associated with shorter work-night sleep and greater free-night rebound, indicating that SJL reflects weekday sleep restriction and compensatory extension of sleep on free nights, in addition to delayed free-night timing. SJL also showed a strong age gradient, with the highest prevalence in younger and middle-aged adults and a marked decline after age 65, likely reflecting earlier free-night sleep timing and closer alignment between work-night and free-night sleep schedules in older patients. Together, these findings suggest that CPAP-derived SJL metrics can provide clinically meaningful information beyond conventional adherence measures by capturing sleep timing, weekly schedule stability, and patterns of weekday restriction with free-night compensation. This information may help clinicians interpret persistent symptoms despite apparently adequate CPAP use and identify patients who may benefit from behavioral sleep counseling, circadian assessment, or further evaluation with sleep diaries or actigraphy.

Overall, CPAP-derived SJL should be viewed as a scalable, low-burden screening phenotype rather than a replacement for dedicated circadian or activity monitoring. Future studies should validate these metrics against actigraphy and sleep diaries, infer individualized work and free-night schedules, quantify within-person stability, and determine whether SJL predicts CPAP adherence, residual symptoms, cardiometabolic risk, or response to behavioral and circadian interventions.

